# Pre-existing H1N1 immunity reduces severe disease with bovine H5N1 influenza virus

**DOI:** 10.1101/2024.10.23.619881

**Authors:** Valerie Le Sage, Bailee D. Werner, Grace A. Merrbach, Sarah E. Petnuch, Aoife K O’Connell, Holly C. Simmons, Kevin R. McCarthy, Douglas S. Reed, Louise H. Moncla, Disha Bhavsar, Florian Krammer, Nicholas A. Crossland, Anita K. McElroy, W. Paul Duprex, Seema S. Lakdawala

## Abstract

The emergence of highly pathogenic H5N1 avian influenza in dairy cattle herds across the United States has caused multiple mild human infections. There is an urgent need to understand the risk of spillover into humans. Here, we show that pre-existing immunity from the 2009 H1N1 pandemic influenza virus provided protection from mortality and severe clinical disease to ferrets intranasally infected with bovine H5N1. H1N1 immune ferrets exhibited a differential tissue tropism with little bovine H5N1 viral dissemination to organs outside the respiratory tract and significantly less H5N1 virus found in nasal secretions and the respiratory tract. Additionally, ferrets with H1N1 prior immunity produced antibodies that cross-reacted with H5N1 neuraminidase protein. Taken together, these results suggest that mild disease in humans may be linked to prior immunity to human seasonal influenza viruses.

## Introduction

In March of 2024, an outbreak of H5N1 clade 2.3.4.4b avian influenza virus was identified in Texas dairy cattle herds and has now spread to over 200 herds in at least 14 states (1). This emphasizes the importance of monitoring this virus for pandemic potential. Infection of various mammals with the 2.3.4.4b clade of H5N1 viruses has resulted in severe disease and mortality in birds, foxes, mink, cats, cetaceans and pinnipeds, but not cows (2, 3). In early April, the first human infection was identified in Texas (4), with a growing number of H5N1 human cases having been identified from workers associated with poultry or dairy farms in California, Missouri, Michigan and Colorado (5). Thus far, human infections in the United States have been characterized by conjunctivitis and mild respiratory symptoms and have not required hospitalizations.

Most individuals experience their first influenza virus infection by the age of five (6), thus current H5N1 human infections are occurring in the presence of prior influenza A virus (IAV) immunity. The reduced disease severity seen in current H5N1 infections could therefore be driven by prior immunity to human seasonal influenza viruses. Statistical modeling analysis of known human cases of H5N1 and H7N9 indicate that childhood HA imprinting may provide lifelong protection against severe infection and death from these viruses (7). Specifically, Gostic *et al*. suggested that immune imprinting with human seasonal H1N1 or H2N2 would reduce disease severity to H5N1 since H5 and H1 and H2 share a similar group 1 HA stalk domain (7). Despite the potential impact of prior immunity to reduce H5N1 replication and pathogenesis, current H5N1 studies have only been performed in immunologically naive ferrets (8). In this work, we report that prior H1N1 immunity reduced virus replication and disease severity of bovine H5N1 virus (clade 2.3.4.4b13) in ferrets. Additionally, we found that ferrets with prior immunity to H1N1 expressed H5N1 cross-reacting antibodies to the neuraminidase protein. Our results suggest that pre-existing immunity to heterotypic influenza viruses may explain the mild symptoms observed so far during H5N1 infection of dairy and poultry farm workers.

## Methods

### Cells

Madin-Darby canine kidney (MDCK) and 293T cells were obtained from American Type Culture Collection (ATCC) and maintained in Minimum Essential medium and Dulbecco’s modified Eagle’s medium (DMEM), respectively. Medium was supplemented with 10 % fetal bovine serum, 2 mM L-glutamine, 100 I.U./mL penicillin and 100 μg/mL streptomycin. All cells were incubated at 37^°^C with 5 % CO2. Human 293F cells were maintained at 37°C with 5-8 % CO2 in FreeStyle 293 Expression Medium (ThermoFisher) supplemented with 100 I.U./mL penicillin and 100 μg/mL streptomycin.

### Rescue of virus from plasmids using co-cultured cells

Reverse genetics plasmids expressing A/dairy cattle/Texas/24008749001/2024 (H5N1) were synthesized based on sequences deposited in the Global Initiative on Sharing All Influenza Data (GIAID) (Accession number EPI_ISL_19014384), with noncoding regions determined from consensus alignment of H5N1 strains from the 2.3.4.4b clade viruses. Each plasmid containing the 8 segments of A/dairy cattle/Texas/24008749001/2024 was diluted to a concentration of 100 ng/ml and a total of 500 ng of each gene segment was combined with Opti-MEM® up to 100 μl and 5 μl of Lipofectamine 2000 transfection reagent (Life Technologies, Waltham, MA). The transfection mixture was incubated at room temperature for 25 min and transferred to 293T cells in Opti-MEM® complete media (Life Technologies, Waltham, MA) in a 6-well plate. After 24 hours of incubation at 37^°^C with 5% CO2, 750,000 MDCK cells were added to the 293T cells. Following another 24-hour incubation, a blind-passage of the rescued virus was performed in MDCK-London cells in a T75 cm^2^ flask. The flask was monitored for cytopathic effect (CPE) for 48 h post-inoculation.

### Virus titration

Nasal wash and organ samples were titered in MDCK cell cultures. Ten-fold serial dilutions were made and inoculated on 96-well plates using 4 wells per dilution. MDCK cells were observed at 4 dpi for cytopathic effect (CPE). Virus titers were calculated using Reed and Muench method (9) and expressed as log10 tissue culture infectious dose 50 (TCID50)/mL.

### Human subjects research ethics statement

The University of Pittsburgh Institutional Review Board approved protocol STUDY20030228 for collection of serum samples from healthy adult donors who provided written informed consent for their samples to be used in infectious disease research. All participants self-reported their age, sex, ethnicity and race.

### Microneutralization assay

The titer of neutralizing antibodies was determined from human or ferret sera that had been heat inactivated at 56^°^C for 30 minutes. Briefly, two-fold serial dilutions of heat-inactivated human serum was incubated with 10^3.3^ TCID50 of influenza virus for 1 hour at room temperature with continuous rocking. Media with tosyl phenylalanyl chloromethyl ketone (TPCK)-treated trypsin was added to 96-well plates with confluent MDCKs before the virus:serum mixture was added. After 4 days, CPE was determined, and the neutralizing antibody titer was expressed as the reciprocal of the highest dilution of serum required to completely neutralize the infectivity of each virus on MDCK cells. The concentration of antibody required to neutralize 100 TCID50 of virus was calculated based on the neutralizing titer dilution divided by the initial dilution factor, multiplied by the antibody concentration.

### Animal ethics statement

Ferret experiments were conducted in BSL2 and BSL3 facilities at the University of Pittsburgh in compliance with the guidelines of the Institutional Animal Care and Use Committee (approved protocol 22061230 and 21089461, respectively). Animals were sedated with isoflurane following approved methods for all nasal washes and survival blood draws. Ketamine and xylazine were used for sedation for all terminal procedures, followed by cardiac administration of euthanasia solution. Approved University of Pittsburgh Division of Laboratory Animal Resources (DLAR) staff administered euthanasia at time of sacrifice. H5N1 studies were performed in accordance with select agent permit number 20230320-074008 (University of Pittsburgh).

### Ferret screening

Four-to six-month-old male ferrets were purchased from Triple F Farms (Sayre, PA, USA). All ferrets were screened by hemagglutinin inhibition (HAI) for antibodies against circulating influenza A and B viruses. The following antigens were obtained through the International Reagent Resource, Influenza Division, WHO Collaborating Center for Surveillance, Epidemiology and Control of Influenza, Centers for Disease Control and Prevention, Atlanta, GA, USA: 2018–2019 WHO Antigen, Influenza A (H3) Control Antigen (A/Singapore/INFIMH-16-0019/2016), β-propiolactone (BPL)-Inactivated, FR-1606; 2014–2015 WHO Antigen, Influenza A (H1N1)pdm09 Control Antigen (A/California/07/2009 NYMC X-179A), BPL-Inactivated, FR-1184; 2018–2019 WHO Antigen, Influenza B Control Antigen, B/Victoria/2/87-like lineage (B/Colorado/06/2017), BPL-Inactivated, FR-1607; 2015–2016 WHO Antigen, Influenza B Control Antigen, B/Yamagata/16/88-like lineage (B/Phuket/3073/2013), BPL-Inactivated, FR-1403.

### Ferret infections

Ferrets with pre-existing immunity against seasonal influenza viruses were experimentally or naturally infected with recombinant A/California/07/2009 (H1N1pdm09). These animals were allowed to recover and housed for >90 days being similarly infected with A/dairy cattle/Texas/24008749001/2024 H5N1 (cow/Tx/24 H5N1). Ferrets with prior H1N1pdm09 immunity or immunologically naive were inoculated intranasally with 10^4^ TCID50 in 500 mL (250 mL in each nostril) with the cow/Tx/24 H5N1 virus. Three animals from each group were euthanized on day 3 for tissue titration and the remaining two were kept for 14 days or until they succumbed to the infection.

### Tissue collection and processing

The respiratory tissues were collected from euthanized ferrets aseptically in the following order: entire right middle lung, left cranial lung (a portion equivalent to the right middle lung lobe), one inch of trachea cut lengthwise, entire soft palate, and nasal turbinates. Tissue samples were weighed, and Leibovitz’s L-15 medium was added to make a 10% (lungs) or 5% (trachea) w/v homogenate. Tissues were dissociated in phosphate-buffered saline (PBS) supplemented with antibiotics and antimycotic using BeadBlaster microtube homogenizer and cell debris was removed by centrifugation at 900 xg for 5 minutes. Influenza virus titers were determined by endpoint TCID50 assay. The lungs were fixed in 10% neutral buffered formalin for two weeks and subsequently processed as formalin fixed paraffin blocks (FFPE) following routine histology processes. Microtomy sections were stained with hematoxylin and eosin (H&E) for histopathologic analysis. Immunohistochemistry (IHC) targeting Influenza A Nucleoprotein (Clone F8L6X, Rb origin; Cell Signaling Technologies) was conducted using a Ventana Discovery Ultra autostainer (Roche, Basel, Switzerland) using a primary concentration of 1:200 and a pre-dilute secondary anti-Rb horseradish peroxidase (HRP) polymer (Vector Laboratories, Newark, California, USA) developed using 3,3’-diaminobenzidine (DAB) chromogen with hematoxylin counterstain (Roche). H&E and IHC slides were scanned using a Phenolmager whole slide scanner (Akoya Biosciences, Malborough, MA, USA) for figure preparation. Slides were initially examined ‘blinded’ to experimental groups to eliminate observer bias by a board-certified veterinary pathologist (NAC), followed by unblinding for figure preparation An ordinal scoring system was developed to summarize the histopathologic and immunohistochemical findings: 0-not observed; 1 (mild), <10% of parenchyma impacted; 2 (moderate) >10%, but <25% of parenchyma impacted; and 3 (severe), >25%, but < 50% of parenchyma impacted. Each of the 5 lung lobes from each ferret were scored individually. Histopathologic features documented included bronchointerstitial pneumonia, perivascular infiltrates, foci of bronchus associated lymphoid tissue (BALT) and influenza A virus nucleoprotein IHC. A cumulative lung injury score was developed encompassing the severity of bronchointerstitial pneumonia and influenza A virus nucleoprotein IHC scores. Perivascular inflammation and foci of BALT were excluded from the cumulative lung injury score as they are interpreted to represent heterotypic adaptive immunity in the cohort with prior exposure to H1N1 prior to H5N1 and thus would falsely elevate lung injury scores if included given the paucity of this phenotype in the cohort without prior influenza A virus immunity to H1N1.

### Recombinant HA expression and purification

Recombinant HA head constructs and full-length HA ectodomains (FLsE) were expressed by polyethylenimine (PEI) facilitated, transient transfection of 293F cells. To produce FLsE constructs, synthetic DNA was subcloned into a pVRC8400 vector encoding a T4 fibritin (foldon) trimerization tag and a 6xHis tag. The H5 dairy cattle HA was modified to contain stabilizing mutations (10) that improved expression and biochemical behavior. Transfection complexes were prepared in Opti-MEM (Gibco) and added to cells. Five days post-transfection, cell supernatants were harvested and clarified by low-speed centrifugation. HA was purified by passage over TALON Metal Affinity Resin (Takara) followed by gel filtration chromatography on Superdex 200 (GE Healthcare) In 10 mM tris(hydroxymethyl)aminomethane (tris), 150 mM NaCl at pH 7.5.

### Recombinant NA expression and purification

Recombinant NA constructs for A/Michigan/45/2015 N1 and A/mallard/New York/22-008760-007-original/2022 N1 were expressed using the baculovirus expression system (11). The constructs were designed to have an N-terminal signal peptide, followed by a hexahistidine purification tag, the VASP (vasodilator-stimulated phosphoprotein) tetramerization domain, a thrombin cleavage site, and the N1 globular head domain. The baculoviruses were passaged in Sf9 cells and then used to infect High Five cells for protein expression. Recombinant proteins were purified 72 hours post-infection from the High Five cell culture supernatant using gravity flow affinity chromatography using Ni^2+^-nitrilotriacetic acid (NTA) agarose (Qiagen).

### ELISA

Five hundred nanograms of rHA FLsE or HA head were adhered to high-capacity binding, 96 well-plates (Corning 9018) overnight in phosphate buffered saline (PBS) pH 7.4 at 4°C. HA or NA coated plates were washed with a PBS-Tween-20 (0.05%v/v) buffer (PBS-T) and then blocked with PBS-T containing 2% bovine serum albumin (BSA) for 1 hour at room temperature. Blocking solution was then removed, and 2-fold dilutions of ferret sera (in blocking solution) were added to wells. Plates were then incubated for 1 hour at room temperature. Primary antibody solution was removed, and plates were washed three times with PBS-T. Secondary antibody, anti-ferret IgG-HRP (Abcam ab97225) diluted 1:10,000 in blocking solution, was added to wells and incubated for 30 minutes at room temperature. Plates were then washed three times with PBS-T. Plates were developed using 150μl 1-Step TMB substrate. Following a brief incubation at room temperature, HRP reactions were stopped by the addition of 100μl of 4N sulfuric acid solution. Plates were read on a Molecular Devices SpectraMax 340PC384 Microplate Reader at 450 nm. All measurements were performed in technical duplicate. The average of the two measurements for each ferret were then graphed as the mean absorbance at 450nm using GraphPad Prism (v9.0).

## Data availability

All source data is available on Figshare (10.6084/m9.figshare.25843414).

## Results

### Low levels of cross-reactive neutralizing antibodies are present in some older individuals

H5N1 influenza A viruses (IAV) have not circulated widely in the human population, and it is unlikely that significant immunity exists against these strains. To assess whether any cross-reactive antibodies existed in the human population, we examined whether sera samples from human donors bracketed by birth year decade could neutralize a H5N1 clade 2.3.4.4b virus strain A/dairy cattle/Texas/24008749001/2024 (cow/Tx/24 H5N1). Neutralization assays conducted with human sera against cow/Tx/24 H5N1 and the 2009 H1N1 pandemic virus (H1N1pdm09) revealed high levels of circulating antibodies against H1N1pdm09 in individuals of all ages, as expected (Figure 1). Surprisingly, 12 of 60 individuals tested had levels of cross-neutralizing antibodies against cow/Tx/24 H5N1 that were above the limit of detection; these individuals were born in the 1940s, 1950s and 1960s, with only two of individuals born after 1970 having detectable cross-neutralizing antibodies (Figure 1), which correlates well with H5 cross-reactive antibodies in older individuals (12). This data indicates that younger individuals could be more susceptible to bovine H5N1 infection. Unfortunately, the ages of the people with documented H5N1 infections since 2022 are not known.

**Figure 1.**
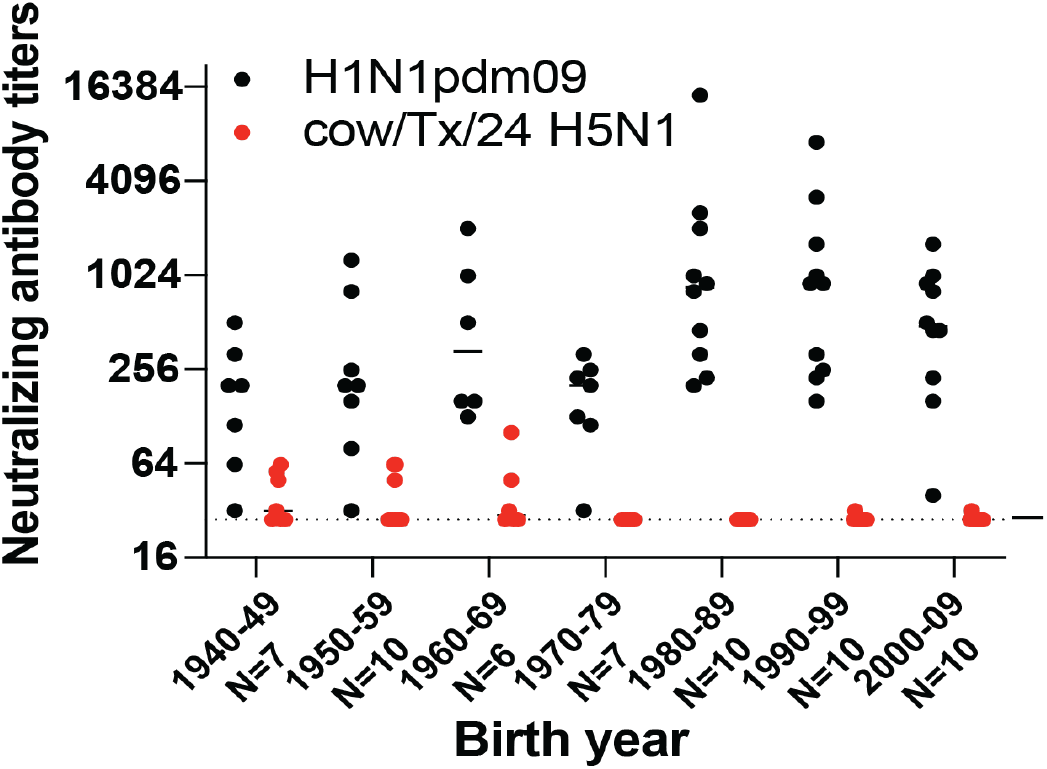
Neutralizing antibody titers of H1N1 and bovine H5N1 in human sera by birth year cohort. Sera collected from the indicated number of healthy individuals in 2020-2021 with birth years ranging from 1940-2009 were tested for neutralizing antibodies against 2009 H1N1 pandemic virus (H1N1pdm09) and cow/Tx/24 H5N1. Each dot represents the neutralizing antibody titer of a single individual to neutralize 100 TCID50 of H1N1pdm09 (black) or cow/Tx/24 H5N1 (red) on MDCK cells. The line indicates the geometric mean value for a given birth decade and the dotted line represents the limit of detection for the assay.

### H1N1pdm09 pre-existing immunity reduces viral titers and dissemination

We previously developed a pre-immune ferret model that has been used to assess the role of prior immunity to human seasonal influenza viruses on infection of heterosubtypic viruses and examine the pandemic potential of circulating swine influenza viruses (13, 14). Prior research has implicated that H1N1 imprinting by birth year could protect from H5N1 infection (7), and therefore, we sought to examine whether this observation would be recapitulated in the ferret model.

To examine the impact of pre-existing immunity on viral replication of bovine H5N1 virus, ferrets were infected with H1N1pdm09 98 days prior to challenge to allow waning of primary immune responses. Ferrets with or without prior H1N1pdm09 immunity were intranasally inoculated with cow/Tx/24 H5N1 virus at a dose of 10^4^ TCID50 and either sacrificed on day 3 (N=3) for assessment of viral load or followed out to day 14 (N=2) to examine mortality (Figure 2A). To examine whether H1N1pdm09 altered cow/Tx/24 H5N1 tissue tropism, intranasally infected ferrets were euthanized on day 3 post-infection to collect tissues (lungs, trachea, soft palate, nasal turbinates, heart, liver, spleen, small intestine, and brain) and virus titers were determined. In ferrets without prior immunity, cow/Tx/24 H5N1 resulted in high viral loads in the respiratory tissues and produced a systemic infection, as observed by virus detection in the heart, liver, spleen, and intestine (Figure 2B). In contrast, ferrets with prior H1N1pdm09 immunity exhibited statistically significant lower levels of virus replication that were limited to the respiratory tract (Figure 2B). The lack of virus in the brain of ferrets with no prior immunity at day 3 is consistent with reported data from other groups (8). Nasal wash titers were also drastically different between the two groups of ferrets. Virus was consistently detected in the nasal washes of ferrets without prior immunity over time, whereas H1N1pdm09 immune ferrets had no detectable cow/Tx/24 H5N1 virus in nasal washes, except for one ferret on day 4 post-infection and a different ferret on day 6 post-infection (Figure 2C).

**Figure 2.**
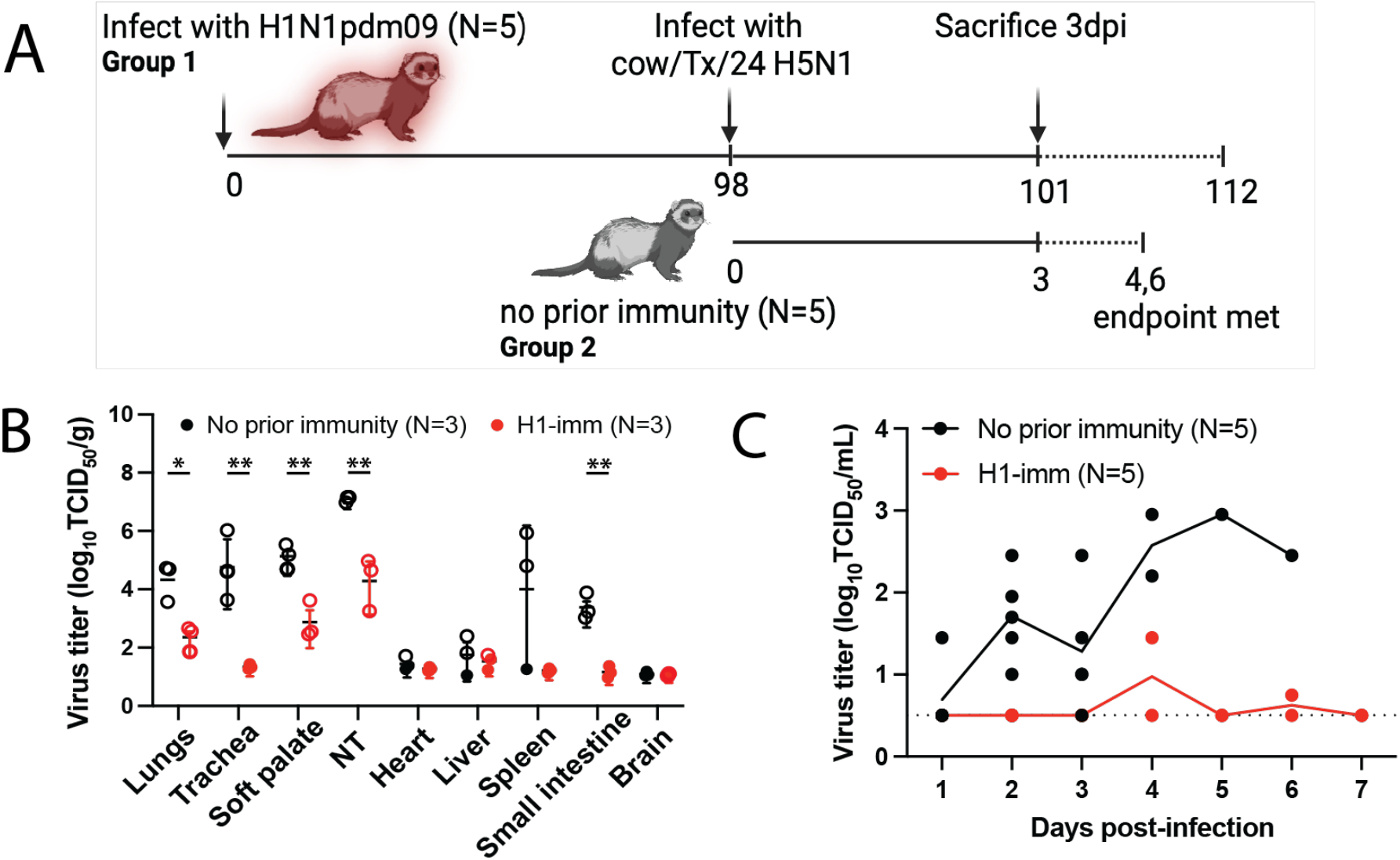
Effects of prior H1N1 immunity on bovine H5N1 virus replication in ferrets. **A**. Schematic of experimental timeline. Two groups of ferrets were intranasally infected with cow/Tx/24 H5N1; group 1 had been infected with H1N1pdm09 98 days prior (N=5) and group 2 were immunologically naïve (N=5). Three animals from each group were sacrificed at day 3 post-infection. The remaining ferrets from group 1 and 2 were monitored until day 14 post-infection or until the endpoint criteria were reached. Schematic was created in BioRender. **B**. Tissues from cow/Tx/24 H5N1 infected ferrets with no prior immunity (black, N=3) or H1N1pdm09 prior immunity (red, N=3) were collected at day 3 post-infection. Mean +/-SD of viral titers are shown with each circle representing an individual ferret. Unpaired t-test analysis was used to determine statistically significant differences (lungs p=0.0124; trachea p<0.0080; soft palate p=0.0072; nasal turbinates (NT) p=0.0061; small intestine p=0.0014). Open circles indicate those values that are above the limit of detection. **C**. Viral titers from nasal secretions of each individual ferret are represented by each circle with a line indicating the mean for each group. Nasal wash samples were collected on the indicated days post-infection (N=5 on days 1-3; N=2 on day 4 until end of the study). The dashed line represents the limit of detection.

Histopathological analysis of lung tissues harvested at day 3 post-infection indicates that although both groups of ferrets present similar lung injury (Figure 3A). However, further refinement of the data indicates that ferrets with pre-existing H1N1pdm09 immunity present with more residual mononuclear perivascular infiltrates and bronchus-associated lymphoid tissue (BALT) hyperplasia (Figure 3B and C), which may play a role in preventing development of severe clinical disease. Immunohistochemistry with anti-IAV nucleoprotein (NP) indicates that H1N1pdm09 immune ferrets had limited NP positive cells in the trachea (Figure 4A) and mainstem bronchi (Figure 4B) and bronchioles (Figure 4C) as compared to ferrets with no prior immunity. IAV NP antigen was observed in alveolar pneumocytes (both type 1 and 2) in ferrets irrespective of immune status (Figure 4C). These findings are in contrast to H1N1 infection in ferrets, where H1N1 virus infects epithelial cells in the large and small airways (15-19). Examination of the tracheobronchial lymph node histology revealed lymphoid depletion, necrosis, fibrin and edema in ferrets with no prior immunity compared to those with H1N1 immunity (Figure 4D). Overall, these data indicate that resident lymphoid changes in ferrets with pre-existing H1N1 immunity may have reduced cow/Tx/24 H5N1 replication and dissemination to other organs, which could impact disease severity.

**Figure 3.**
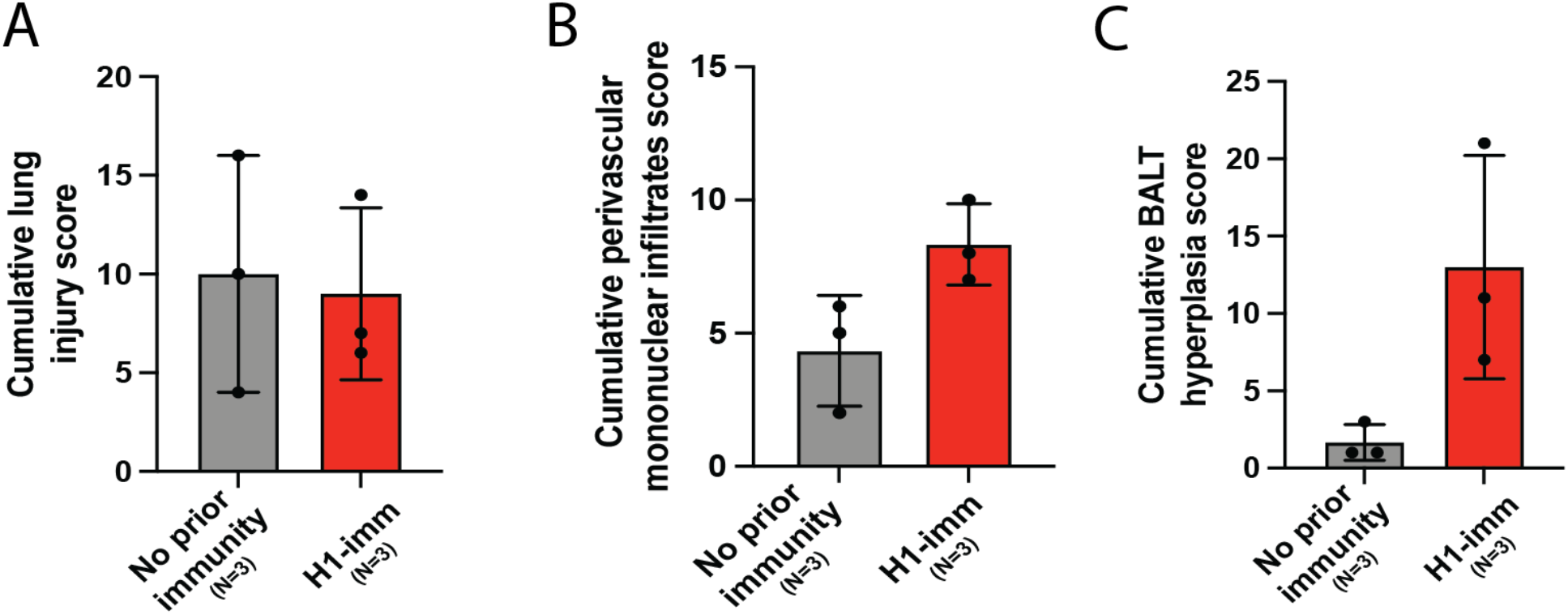
Pre-existing H1N1 immunity increases lung immune infiltrates. Five lung sections for ferrets with no prior (gray) or pre-existing H1N1pdm09 (red) immunity were blindly scored for **A**. lung injury, **B**. perivascular mononuclear infiltrates and **C**. BALT hyperplasia. Each dot represents the cumulative score of each of the five sections for each ferret.

**Figure 4.**
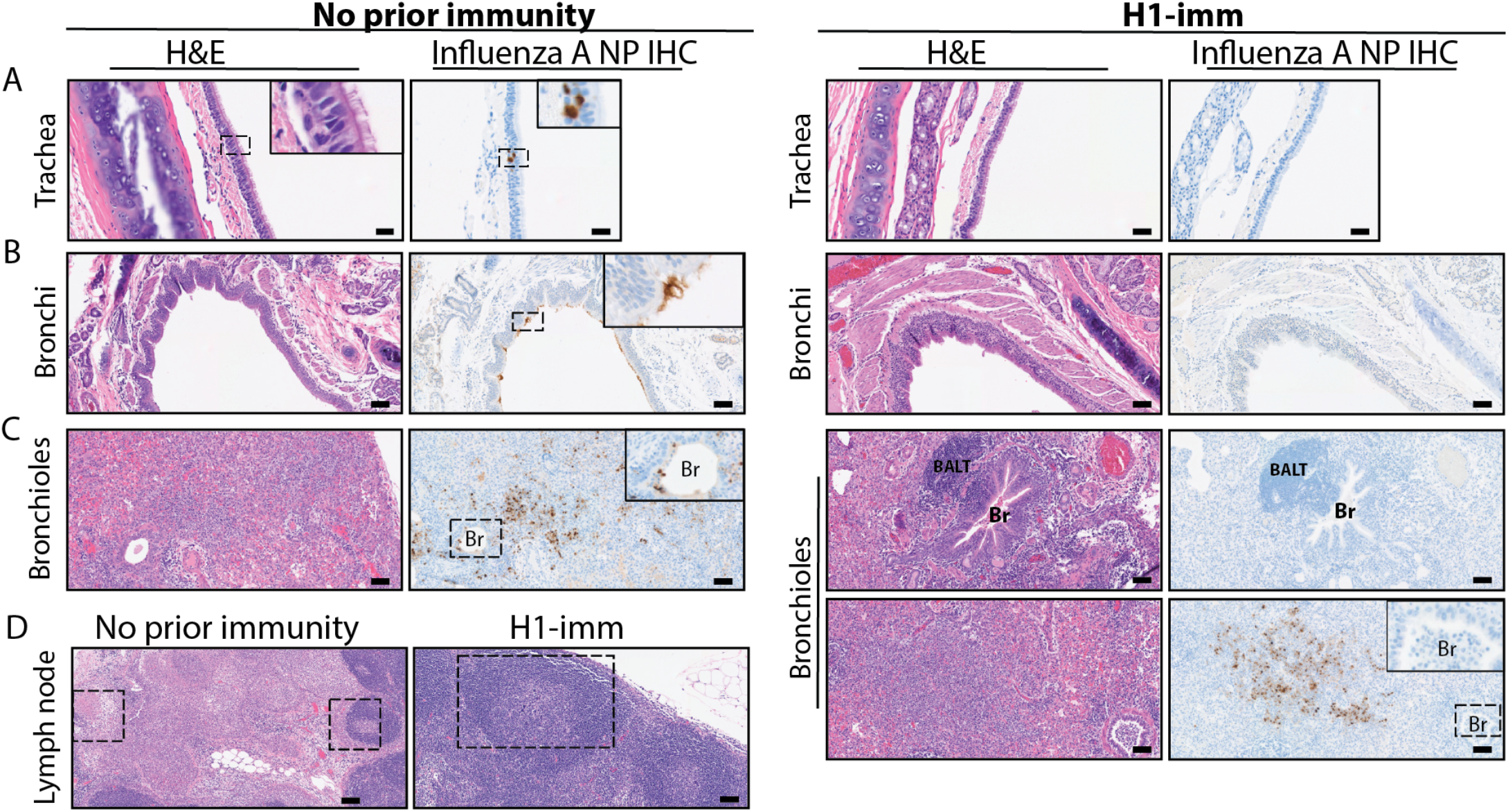
Pre-existing H1N1 immunity reduces the presence of viral antigen and increases immune infiltrates. Ferrets with no prior immunity (left panels) or H1N1pdm09 pre-existing immunity (right panels) were infected with 10^4^ TCID50 of cow/Tx/24 H5N1 and sacrificed on day 3 post-infection. H&E and influenza A virus nucleoprotein IHC were performed for the trachea (**A**, images taken at 400x with 20mm scale bars), bronchi (**B**, all images taken at 200x with 50mm scale bars) and bronchiole (BR) (**C**, all images taken at 200x with 50mm scale bars). BALT refers to bronchus-associated lymphoid tissue. Hatched squares are areas within parts A-C that are magnified within the inset panel. **D**. H&E of tracheobronchial lymph node from ferrets with no prior immunity (left panel) or H1N1pdm09 pre-existing immunity (right panel) (images taken at 100x with 100mm scale bars, insets taken at 400x with 20mm scale bars). Hatched squares in part D indicate areas with immune infiltration.

### H1N1pdm09 immunity protects against mortality and severe disease

Ferrets with or without prior H1N1pdm09 immunity were followed out to day 14 (N=2) to examine mortality (Figure 5A). All animals with pre-existing H1N1pdm09 immunity survived challenge with the cow/Tx/24 H5N1 virus, while immunologically naive ferrets succumbed to the infection on days 4 and 6 post-infection (Figure 5A). No more than 5% weight loss was observed in ferrets with pre-existing H1N1pdm09 immunity (Figure 5B) and further assessment of clinical signs, such as diarrhea, fever, nasal discharge and playfulness, revealed more severe clinical signs in all immunologically naive animals compared to those with prior H1N1pdm09 immunity (Figure 5C). Surviving ferrets with H1N1pdm09 pre-existing immunity seroconverted against cow/Tx/24 H5N1, albeit to low microneutralization titers of 20 and 80 (Table 1). Additionally, neither of the two H1N1pdm09 immune ferrets had a greater than 4-fold rise in anti-H1N1pdm09 antibodies post-cow/Tx/24 H5N1 challenge (Table 1). Taken together, these data indicate that H1N1pdm09 pre-existing immunity protects ferrets from severe clinical disease and mortality caused by cow/Tx/24 H5N1 infection.

**Table 1:** Serology of cow/Tx/24 H5N1 infected ferrets.

| Ferret | Immunity | Sacrifice dpi | cow/Tx/24 MN titers (D0) | cow/Tx/24 MN titers (D3/4/6) | H1N1pdm09 MN titers (D0) | H1N1pdm09 MN titers (D3/14) |
| --- | --- | --- | --- | --- | --- | --- |
| 1 | No prior | 3 | <20 | <20 | ND | ND |
| 2 | No prior | 3 | <20 | <20 | ND | ND |
| 3 | No prior | 3 | <20 | <20 | ND | ND |
| 4 | No prior | 4 | <20 | <20 | ND | ND |
| 5 | No prior | 6 | <20 | <20 | ND | ND |
| 1 | H1-imm | 3 | <20 | <20 | 905 | 1016 |
| 2 | H1-imm | 3 | <20 | <20 | 508 | 320 |
| 3 | H1-imm | 3 | <20 | <20 | 2032 | 508 |
| 4 | H1-imm | 14 | <20 | 20 | 2560 | 2032 |
| 5 | H1-imm | 14 | <20 | 80 | 640 | 2032 |
| ND = Not determined |  |  |  |  |  |  |
| MN = microneutralization |  |  |  |  |  |  |

MN = microneutralization
MN = microneutralization

**Figure 5.**
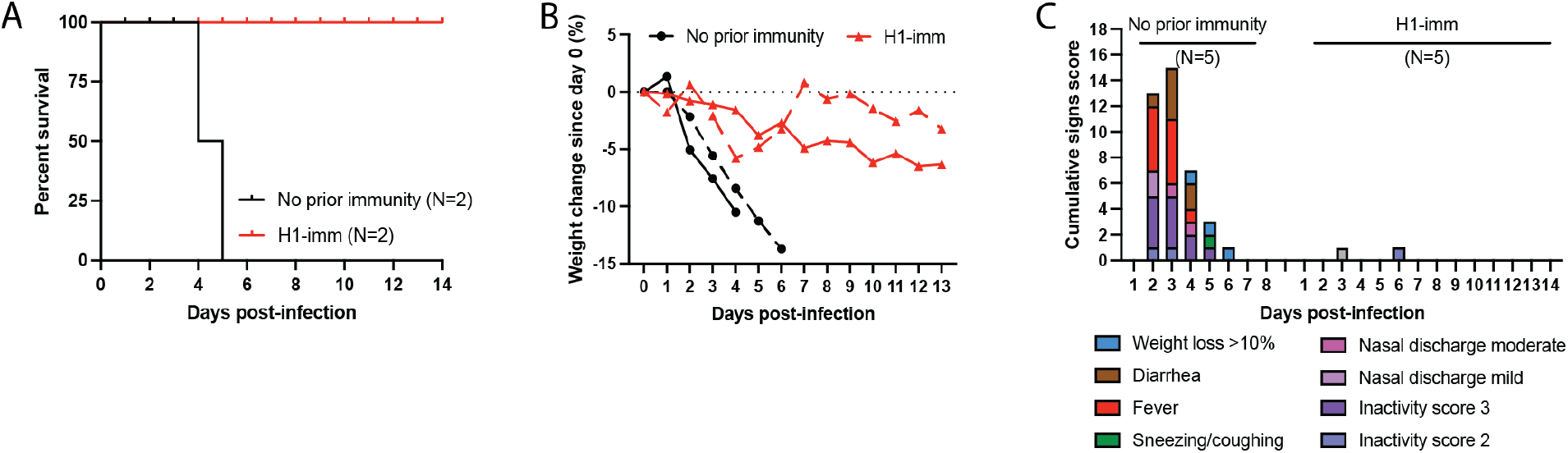
Prior H1N1 immunity protects from mortality and severe disease in ferrets infected with bovine H5N1 virus. **A**. Mortality of ferrets in each group (N=2 per group), with no prior immunity shown in black and H1N1pdm09 pre-existing immunity (H1-imm) in red. **B**. Percent of weight change for ferrets with no prior immunity (black, N=2) and H1N1pdm09 pre-existing immunity (H1-imm, red, N=2). **C**. Clinical signs on the indicated days post-infection (N=5 on days 1-3; N=2 on day 4 until end of the study) for ferrets with no prior immunity or H1N1pdm09 pre-existing immunity (H1-imm). Symptoms of infection were monitored each day post-infection and quantified into a cumulative signs score.

### H1N1 immune ferrets have cross-reactive NA antibodies prior to challenge with H5N1

IAV infection induces antibody responses against the HA and NA that can provide varying levels of protection against subsequent infections (20). To identify immune factors that contribute to the protection of H1N1 immune ferrets from severe disease, neutralizing and total HA binding antibodies were measured. Before challenge with cow/Tx/24 H5N1 virus, ferrets with H1N1 prior immunity exhibited high levels of neutralizing antibodies against H1N1pdm09 but no neutralizing antibodies above the limit of detection against cow/Tx/24 H5N1 (Figure 6A). Cross-reactive HA stalk-specific antibodies are able to play a role in reducing influenza virus disease severity (21-23). To explore the production of non-neutralizing cross-reactive HA antibodies, an enzyme-linked immunosorbent assay (ELISA) using the whole H1 (A/California/07/2009 H1N1) or H5 (A/dairy cattle/Texas/24008749001/2024 H5N1) HA protein was performed with serum from ferrets with pre-existing H1N1pdm09 immunity (Figure 6B). Ferrets with H1N1pdm09 prior immunity produced antibodies that bound to H1 as expected but displayed the same background levels of antibody binding to the H5 HA protein as ferrets with no prior immunity (Figure 6B), indicating that there are no detectable cross-reactive HA antibodies against the avian H5. Finally, an ELISA was performed using neuraminidase (NA) from a human (A/Michigan/45/2015 H1N1) or avian (A/mallard/New York/22-008760-007-original/2022 H5N1; which is 98.7% similar to cow/Tx/24 NA) IAV to determine whether H1N1 pre-immune ferrets had any cross-reacting NA antibodies prior to challenge with H5N1 that might contribute to the protection against severe disease. Surprisingly, sera from ferrets with prior H1N1pdm09 immunity had antibodies that bound to both the human and avian NA antigens, while the ferrets with no prior immunity had background levels of binding (Figure 6C). These data suggest that cross-reactive NA antibodies to the avian N1 may be produced from a human seasonal H1N1 infection.

**Figure 6.**
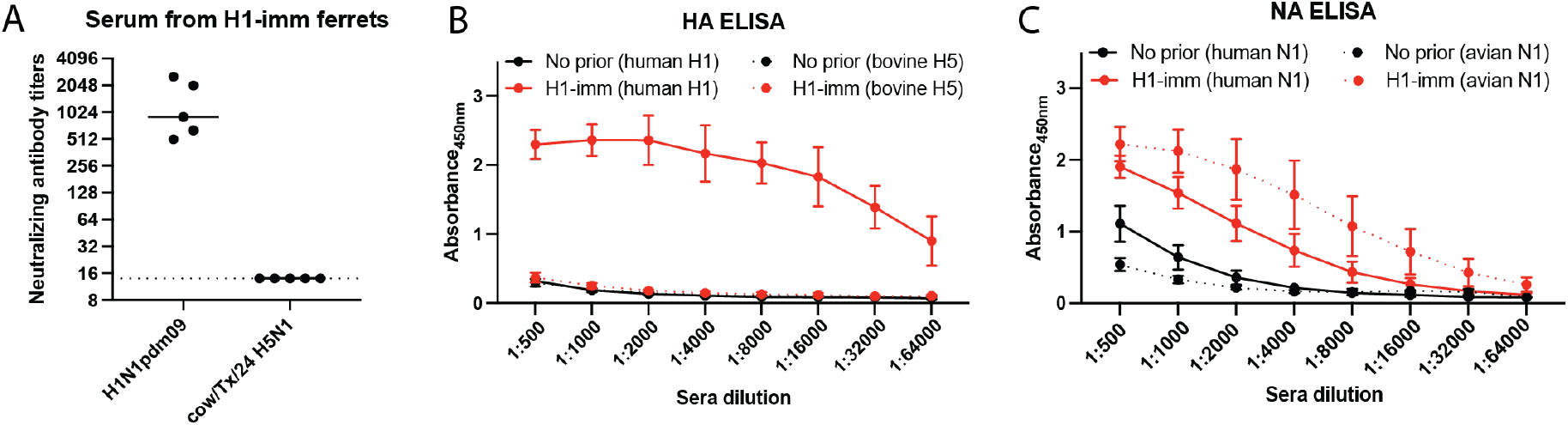
Ferrets with H1N1pdm09 pre-existing immunity have cross-reactive NA binding antibodies on day of challenge. **A**. Sera collected from the five ferrets with pre-existing H1N1pdm09 immunity on day 98 post-infection were tested for neutralizing antibodies against 2009 H1N1 pandemic virus (H1N1pdm09) and cow/Tx/24 H5N1. Each dot represents the neutralizing antibody titer of a single ferret to neutralize 100 TCID50 of H1N1pdm09 or cow/Tx/24 H5N1 on MDCK cells. The line indicates the geometric mean value for each virus and the dotted line represents the limit of detection for the assay. **B**. Serum IgG antibodies in ferrets with no prior (black) or with pre-existing H1N1pdm09 immunity (red) against purified HA proteins. The solid lines show ferret serum reactivity to human HA (A/Michigan/45/2015 H1N1) and the dashed lines show ferret serum reactivity to bovine HA (A/dairy cattle/ Texas/24008749001/2024 H5N1). Data is presented as mean +/-SD of the absorbance at 450nm for each dilution. **C**. Serum IgG antibodies in ferrets with no prior (black) or with pre-existing H1N1pdm09 immunity (red) against purified NA proteins. The solid lines show ferret serum reactivity to human NA (A/California/07/2009 H1N1) and the dashed lines show ferret serum reactivity to avian NA (A/mallard/New York/22-008760-007-original/2022 H5N1). Data is presented as mean +/-SD of the absorbance at 450nm for each dilution.

## Discussion

H1N1pdm09 pre-existing immunity in ferrets was sufficient to protect from severe disease and mortality from the highly pathogenic avian influenza bovine H5N1 virus. Significantly reduced H5N1 viral titers in nasal secretions and respiratory tract were also observed in the animals with H1N1 immunity. Importantly, protection from H5N1 infection was not due to the presence of cross-neutralizing antibodies in sera as ferrets with H1N1pdm09 immunity did not generate systemic antibodies that cross-neutralized the cow/Tx/24 H5N1 virus (Figure 6A). Rather, ferrets with prior immunity to H1N1 were found to produce cross-reacting antibodies to H5N1 NA protein (Figure 6C); this observation is consistent with recently reported human serological data (24). Immunity to NA has previously been implicated to provide protection during the 1968 H3N2 pandemic (25, 26) and can reduce disease severity of naturally infected individuals and those experimental challenge (27).

While anti-NA antibodies may be facilitating the protection from severe disease observed in the H1N1pdm09 immune ferret, further studies on the mechanisms of protection are clearly warranted and should include an examination of mucosal immunity from antibodies in the respiratory tract that have broad binding potential. Tissue-resident memory T-cells may also help reduce the severity of disease, as is suspected in the case of H1 immunity protecting from airborne transmission of human seasonal H3N2 virus (13). A conservation of immunodominant T cell epitopes between H5N1 and seasonal influenza viruses, including H1N1, was recently reported and suggested to potentially provide a level of cross-protective immunity (28). We did note that the lung tissues of ferrets with H1N1 prior immunity had increased mononuclear perivascular infiltrates and bronchus-associated lymphoid tissue hyperplasia, consistent with tissue-specific T cell responses, although additional investigation is required.

The mild infection that was present in the two H1N1pdm09-immune ferrets that survived until day 14, might account for the low levels of neutralizing antibodies against cow/Tx/24 H5N1, (Table 1). This observation may be critical to inform the use of H5 seroconversion as a detection mechanism for prevalence of H5 infections in farm workers, since mild infections may not produce a robust systemic antibody immune response.

All adults have pre-existing immunity from repeated influenza virus infections over their lifetime. Most of the 2022 and onward H5N1 human case reports have not included the age of the dairy and poultry farm workers infected. However, it is likely they are younger than 50 or 60 years of age, and thus would be highly susceptible to H5N1 infection, yet have circulating H1N1 influenza antibodies. The disease presentation of the current H5N1 human cases is in stark contrast to those reported in the early 2000’s where 30-50% mortality was observed (29). The difference in disease presentation could be due to a number of factors, including changes in the viral genome that result in a less pathogenic virus or the impact of prior immunity to H1N1 strains that circulate widely post 2010. However, additional research into the level of protection afforded by other human seasonal influenza viruses, particularly currently circulating H1N1 viruses and those prior to the 2009 H1N1 pandemic, are needed to assess whether currently circulating H1N1 viruses produce a protective immune signature while other prior strains did not.

## First Author Biography

Dr. Le Sage is a Research Assistant Professor at the University of Pittsburgh Center for Vaccine Research. Her research interests include elucidating the requirements for influenza virus transmission and assessing the pandemic potential of emerging influenza viruses.

## Acknowledgements

This project has been funded in part with Federal funds from the National Institute of Allergy and Infectious Diseases, National Institutes of Health, Department of Health and Human Services, under Contract No. 75N93021C00015 and an NIH award 5R01AI58484-03 to SSL. An NIH award (UC7AI180311) from the National Institute of Allergy and Infectious Diseases (NIAID) supporting the Operations of The University of Pittsburgh Regional Biocontainment Laboratory (RBL) within the Center for Vaccine Research (CVR). Two NIH S10 instrumentation awards (S10OD030269 & S10OD026983) to NAC. We would like to extend our deepest gratitude to Dr. Julian Arthur and Jessica Simendinger, at Cell Signaling Technology, Inc., for their outstanding work in developing the Influenza A Nucleoprotein (NP) (F8L6X) Rabbit mAb #99797. Their expertise, dedication, and tireless efforts have been instrumental in the successful creation of this critical clone, and we are particularly grateful for their collaborative efforts with our team during the validation process. Burroughs Wellcome CAMS 1013362.02 to AKM. We thank Dr. Rachel Duron for critical review and feedback. Work in the Krammer laboratory was supported via an NIAID-funded Center of Excellence for Influenza Research and Response (CEIRR, contract number 75N93021C00014).

## Author contributions

VL and SSL designed the experiments, analyzed, interpreted the data and wrote the manuscript. VL, BDW, GAM, SEP, AKO, HCS and NAC performed the experiments. KRM, DSR, LHM, DB, FK, AKM, and WPD contributed resources and analysis. All authors edited and approved the manuscript.

## References

1. HPAI Confirmed Cases in Livestock 2024 [Available from: https://www.aphis.usda.gov/livestock-poultry-disease/avian/avian-influenza/hpai-detections/hpai-confirmed-cases-livestock.

2. Koopmans MPG, Barton Behravesh C, Cunningham AA, Adisasmito WB, Almuhairi S, Bilivogui P, et al. The panzootic spread of highly pathogenic avian influenza H5N1 sublineage 2.3.4.4b: a critical appraisal of One Health preparedness and prevention. Lancet Infect Dis. 2024.

3. Peacock T, Moncla L, Dudas G, VanInsberghe D, Sukhova K, Lloyd-Smith JO, et al. The global H5N1 influenza panzootic in mammals. Nature. 2024.

4. Uyeki TM, Milton S, Abdul Hamid C, Reinoso Webb C, Presley SM, Shetty V, et al. Highly Pathogenic Avian Influenza A(H5N1) Virus Infection in a Dairy Farm Worker. N Engl J Med. 2024;390(21):2028–9.

5. CDC. Avian Influenza (Bird Flu) [Available from: https://www.cdc.gov/bird-flu/spotlights/ah5n1-response-update.html.

6. Bodewes R, de Mutsert G, van der Klis FR, Ventresca M, Wilks S, Smith DJ, et al. Prevalence of antibodies against seasonal influenza A and B viruses in children in Netherlands. Clin Vaccine Immunol. 2011;18(3):469–76.

7. Gostic KM, Ambrose M, Worobey M, Lloyd-Smith JO. Potent protection against H5N1 and H7N9 influenza via childhood hemagglutinin imprinting. Science. 2016;354(6313):722–6.

8. Eisfeld AJ, Biswas A, Guan L, Gu C, Maemura T, Trifkovic S, et al. Pathogenicity and transmissibility of bovine H5N1 influenza virus. Nature. 2024.

9. Reed LJaM, H. A simple method of estimating fifty per cent endpoints. Am J Epidemiol. 1938;27:493–7.

10. Milder FJ, Jongeneelen M, Ritschel T, Bouchier P, Bisschop IJM, de Man M, et al. Universal stabilization of the influenza hemagglutinin by structure-based redesign of the pH switch regions. Proc Natl Acad Sci U S A. 2022;119(6).

11. Margine I, Palese P, Krammer F. Expression of functional recombinant hemagglutinin and neuraminidase proteins from the novel H7N9 influenza virus using the baculovirus expression system. J Vis Exp. 2013(81):e51112.

12. Nachbagauer R, Choi A, Izikson R, Cox MM, Palese P, Krammer F. Age Dependence and Isotype Specificity of Influenza Virus Hemagglutinin Stalk-Reactive Antibodies in Humans. mBio. 2016;7(1):e01996–15.

13. Le Sage V, Jones JE, Kormuth KA, Fitzsimmons WJ, Nturibi E, Padovani GH, et al. Pre-existing heterosubtypic immunity provides a barrier to airborne transmission of influenza viruses. PLoS Pathog. 2021;17(2):e1009273.

14. Le Sage V, Rockey NC, French AJ, McBride R, McCarthy KR, Rigatti LH, et al. Potential pandemic risk of circulating swine H1N2 influenza viruses. Nat Commun. 2024;15(1):5025.

15. Camp JV, Bagci U, Chu YK, Squier B, Fraig M, Uriarte SM, et al. Lower Respiratory Tract Infection of the Ferret by 2009 H1N1 Pandemic Influenza A Virus Triggers Biphasic, Systemic, and Local Recruitment of Neutrophils. J Virol. 2015;89(17):8733–48.

16. Lakdawala SS, Jayaraman A, Halpin RA, Lamirande EW, Shih AR, Stockwell TB, et al. The soft palate is an important site of adaptation for transmissible influenza viruses. Nature. 2015;526(7571):122–5.

17. Lakdawala SS, Shih AR, Jayaraman A, Lamirande EW, Moore I, Paskel M, et al. Receptor specificity does not affect replication or virulence of the 2009 pandemic H1N1 influenza virus in mice and ferrets. Virology. 2013;446(1-2):349–56.

18. Smith JH, Nagy T, Driskell E, Brooks P, Tompkins SM, Tripp RA. Comparative pathology in ferrets infected with H1N1 influenza A viruses isolated from different hosts. J Virol. 2011;85(15):7572–81.

19. Vidana B, Martinez J, Martinez-Orellana P, Garcia Migura L, Montoya M, Martorell J, et al. Heterogeneous pathological outcomes after experimental pH1N1 influenza infection in ferrets correlate with viral replication and host immune responses in the lung. Vet Res. 2014;45(1):85.

20. Kosik I, Yewdell JW. Influenza Hemagglutinin and Neuraminidase: Yin(-)Yang Proteins Coevolving to Thwart Immunity. Viruses. 2019;11(4).

21. Pica N, Hai R, Krammer F, Wang TT, Maamary J, Eggink D, et al. Hemagglutinin stalk antibodies elicited by the 2009 pandemic influenza virus as a mechanism for the extinction of seasonal H1N1 viruses. Proc Natl Acad Sci U S A. 2012;109(7):2573–8.

22. Throsby M, van den Brink E, Jongeneelen M, Poon LL, Alard P, Cornelissen L, et al. Heterosubtypic neutralizing monoclonal antibodies cross-protective against H5N1 and H1N1 recovered from human IgM+ memory B cells. PLoS One. 2008;3(12):e3942.

23. Wu NC, Wilson IA. Influenza Hemagglutinin Structures and Antibody Recognition. Cold Spring Harb Perspect Med. 2020;10(8).

24. Daulagala P, Cheng SMS, Chin A, Luk LLH, Leung K, Wu JT, et al. Avian Influenza A(H5N1) Neuraminidase Inhibition Antibodies in Healthy Adults after Exposure to Influenza A(H1N1)pdm09. Emerg Infect Dis. 2024;30(1):168–71.

25. Monto AS, Kendal AP. Effect of neuraminidase antibody on Hong Kong influenza. Lancet. 1973;1(7804):623–5.

26. Murphy BR, Kasel JA, Chanock RM. Association of serum anti-neuraminidase antibody with resistance to influenza in man. N Engl J Med. 1972;286(25):1329–32.

27. Maier HE, Nachbagauer R, Kuan G, Ng S, Lopez R, Sanchez N, et al. Pre-existing Antineuraminidase Antibodies Are Associated With Shortened Duration of Influenza A(H1N1)pdm Virus Shedding and Illness in Naturally Infected Adults. Clin Infect Dis. 2020;70(11):2290–7.

28. Sidney J, Kim A-R, de Vries RD, Peters B, Meade PS, Krammer F, et al. Targets of influenza Human T cell response are mostly conserved in H5N1. bioRxiv. 2024:2024.09.09.612060.

29. Organization WH. Cumulative number of confirmed human cases for avian influenza A(H5N1) reported to WHO, 2003-2024, 26 February 2024 [Available from: https://www.who.int/publications/m/item/cumulative-number-of-confirmed-human-cases-for-avian-influenza-a(h5n1)-reported-to-who--2003-2024-26-february-2024.

